# Text mining of 15 million full-text scientific articles

**DOI:** 10.1101/162099

**Authors:** David Westergaard, Hans-Henrik Stærfeldt, Christian Tønsberg, Lars Juhl Jensen, Søren Brunak

**Affiliations:** Center for Biological Sequence Analysis, Department of Bio and Health Informatics, Technical University of Denmark, DK-2800 Lyngby, Denmark; Novo Nordisk Foundation Center for Protein Research, Faculty of Health and Medical Sciences, University of Copenhagen, DK-2200 Copenhagen, Denmark; Office for Innovation and Sector Services, Technical Information Center of Denmark, Technical University of Denmark, DK-2800 Lyngby, Denmark

## Abstract

Across academia and industry, text mining has become a popular strategy for keeping up with the rapid growth of the scientific literature. Text mining of the scientific literature has mostly been carried out on collections of abstracts, due to their availability. Here we present an analysis of 15 million English scientific full-text articles published during the period 1823–2016. We describe the development in article length and publication sub-topics during these nearly 250 years. We showcase the potential of text mining by extracting published protein–protein, disease–gene, and protein subcellular associations using a named entity recognition system, and quantitatively report on their accuracy using gold standard benchmark data sets. We subsequently compare the findings to corresponding results obtained on 16.5 million abstracts included in MEDLINE and show that text mining of full-text articles consistently outperforms using abstracts only.

## Introduction

Text mining has become a widespread approach to identify and extract information from unstructured text. Text mining is used to extract facts and relationships in a structured form that can be used to annotate specialized databases, to transfer knowledge between domains and more generally within business intelligence to support operational and strategic decision-making [1–3]. Biomedical text mining is concerned with the extraction of information regarding biological entities, such as genes and proteins, phenotypes, or even more broadly biological pathways (reviewed extensively in [3–9]) from sources like scientific literature, electronic patient records, and most recently patents [10–13]. Furthermore, the extracted information has been used as annotation of specialized databases and tools (reviewed in [3,14]). In addition, text mining is routinely used to support manual curation of biological databases [15,16]. Thus, text mining has become an integral part of many resources serving a wide audience of scientists. The main text source for scientific literature has been the MEDLINE corpus of abstracts, essentially due to the restricted availability of full-text articles. However, full-text articles are becoming more accessible and there is a growing interest in text mining of complete articles. Nevertheless, to date no studies have presented a systematic comparison of the performance comparing abstracts and full-texts in corpora that are similar in size to MEDLINE.

Full-text articles and abstracts are structurally different [17]. Abstracts are comprised of shorter sentences and very succinct text presenting only the most important findings. By comparison, full-text articles contain complex tables, display items and references. Moreover, they present existing and generally accepted knowledge in the introduction (often presented in the context of summaries of the findings), and move on to reporting more in-depth results, while discussion sections put the results in perspective and mention limitations and concerns. The latter is often considered to be more speculative compared to the abstract [3].

While text-mining results from accessible full-text articles have already become an integral part of some databases (reviewed recently for protein-protein intaractions [18]), very few studies to date have compared text mining of abstracts and full-text articles. Using a corpus consisting of ~20,000 articles from the PubMed Central (PMC) open-access subset and Directory of Open Access Journals (DOAJ), it was found that many explicit protein–protein interactions only are mentioned in the full text [19]. Additionally, in a corpus of 1,025 full-text articles it was noticed that some pharmacogenomics associations are only found in the full text [20]. One study using a corpus of 3,800 articles with focus on *Caenorhabditis elegans* noted an increase in recall from 45% to 95% when including the full text [21]. Other studies have worked with even smaller corpora [17,22,23]. One study have even noted that the majority of claims within an article is not reported in the abstract [24]. Whilst these studies have been of significant interest, the number of full-text articles and abstracts used for comparison are nowhere near the magnitude of the actual number of scientific articles published to date, and it is thus unclear if the results can be generalized to the scientific literature as a whole. The earlier studies have mostly used articles retrieved from PMC in a structured XML file. However, full-text articles received or downloaded directly from the publishers often come in the PDF format, which must be converted to a raw unformatted text file. This presents a challenge, as the quality of the text mining will depend on the proper extraction and filtering of the unformatted text. A previous study dealt with this by writing custom software taking into account the structure and font of each journal at that time [21]. More recent studies typically provide algorithms that automatically determines the layout of the articles [25–27].

In this work, we describe a corpus of 15 million full-text scientific articles from Elsevier, Springer, and the open-access subset of PMC. The articles were published during the period 1823–2016. We highlight the possibilities by extracting protein–protein associations, disease–gene associations, and protein subcellular localization from the large collection of full-text articles using a Named Entity Recognition (NER) system combined with a scoring of co-mentions. We quantitatively report the accuracy and performance using gold standard benchmark data sets. Lastly, we compare the findings to corresponding results obtained on the matching set of abstracts included in MEDLINE as well as the full set of 16.5 million MEDLINE abstracts.

## Results

### Growth and temporal development in full text corpora

The growth of the data set over time is of general interest in itself, however, it is also important to secure that the concepts used in the benchmarks are likely to be present in a large part of the corpus. We found that the number of full-text articles has grown exponentially over a long period (Fig 1a, a log-transformed version is provided in Supplementary Fig 1). We also observed that the growth represents a mixture of two components: one from 1823–1944, and another from 1945–2016. Fitting an exponential curve to the years 1945–2016 we found that the growth rate is 0.103 (p < 2 * 10^−16^, R^2^ = 0.95). Thus, the doubling time for the full-text corpus is 9.7 years. In comparison, MEDLINE had a growth rate of 0.195 (p < 2 * 10^−16^, R^2^ = 0.91) and a doubling time of 5.1 years. We noticed that there was a drop in the number of full-text publications around the years 1914–1918 and 1940–1945. Likewise, we see a decrease in the number of publications indexed by MEDLINE in the entire period 1930–1948.

**Fig 1:**
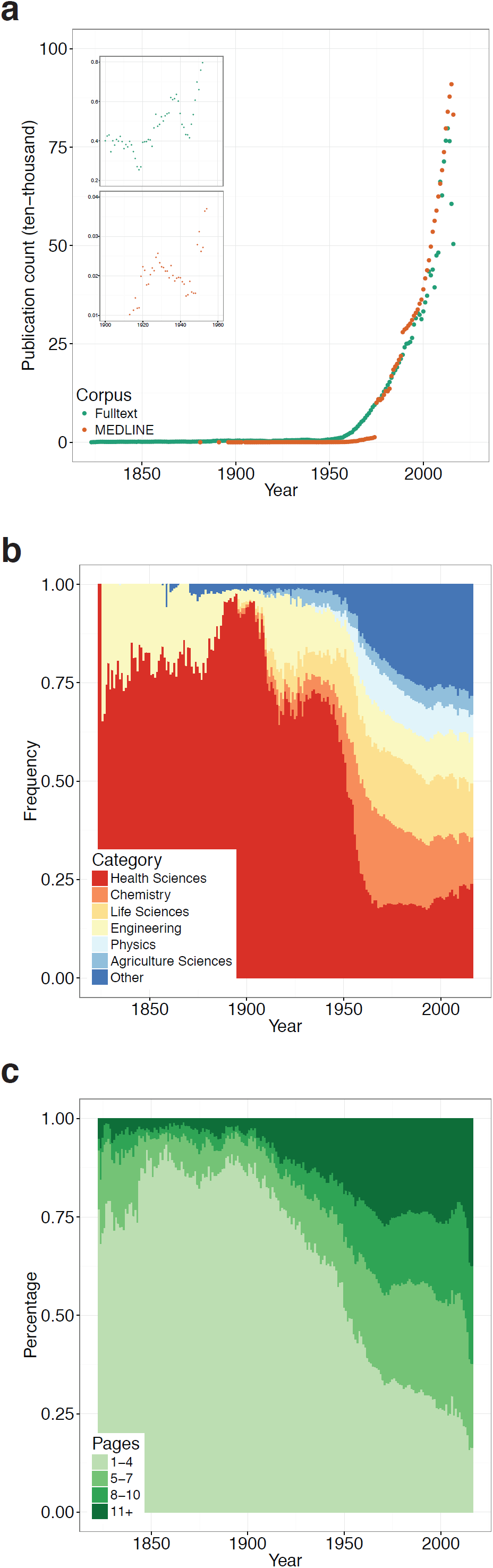
Temporal corpus statistics. **(a)** Number of publications per year in the period 1823-2016. The growth in publications was found to fit an exponential model. **(b)** Temporal development in the distribution of six different topical categories in the period 1823-2016. Publications from health science journals made up nearly 75% of all publications until 1950, at which point it started to decrease rapidly. To date, it makes up approximately 25% of the publications in the full-text corpus. **(c)** Development in the number of pages per article in the period 1823-2016. The range of pages varies from 1-1,572 pages. Until year 1900 the number of one-page articles were increasing, at one point making up 75% of all articles. At the end of the 19th century, the number of one-page articles started to decrease, and by the start of the 21th century they made up less than 20%. Conversely, the number of articles with 11+ pages has been increasing, and by the start of the 21th century made up more than 20% of all articles.

We binned the full-text articles into four categories based on the number of pages (see Methods). The average length of articles has increased considerably during the almost 250 years studied (Fig 1b). Whereas 75% of the articles were 1–3 pages long at the end of the 20^th^ century, less than 25% of the articles published after year 2000 are that short. Conversely, articles with ten or more pages only made up between 0.7%-7% in the 19^th^ century, a level that had grown to 20% by the start of the 21^st^ century.

In the full-text corpora we found a total of 12,781 unique journal titles. The most prevalent journals are tied to health or life sciences, such as *The Lancet, Tetrahedron Letters,* and *Biochemical and Biophysical Research Communications,* or the more broad journals such as *PLoS ONE* (see Supplementary Table 1 for the top-15 journals). *The Lancet* publishes only very few articles per issue, it was established in 1823 and has been active in publishing since then, thus explaining why it so far has nearly published 400,000 articles. In contrast, *PLoS ONE* was launched in 2006, and has published more than 172,000 articles. Of the 12,781 journal titles, 6,900 had one or more category labels assigned by librarians at the Technical University of Denmark. The vast majority of the full-texts, 13,343,040, were published in journals with one or more category labels. The frequency of each category within the corpus can be seen in Supplementary Fig 2. We observed that before the 1950’s health science dominated and made up almost 75% of all publications (Fig 1c). At the start of the 1950’s the fraction started to decrease, and to date health science makes up approximately 25% of all publications in the full-text corpus. Inspecting the remaining eleven categories in a separate plot we found that there was no single category that was responsible for the growth (Supplementary Fig 3).

### Evaluating information extraction across corpora

We analyzed and compared four different corpora comprising all full-text articles (14,549,483 articles, All Full-texts), full-text articles that had a separate abstract (10,376,626 articles, Core Full-texts), the abstract from the full-text articles (10,376,626 abstracts, Core Abstracts), and the MEDLINE corpus (16,544,511 abstracts, MEDLINE). We have used quite difficult, but still well established benchmarks, to illustrate the differences in performance when comparing text mining of abstracts to full-text articles. Within biology, and specifically in the area of systems biology, macromolecular interactions and the relationships between genes, tissues and diseases are key data that drive modeling and the analysis of causal biochemical mechanisms. Knowledge of interactions between proteins is extremely useful when revealing the components, which contribute to mechanisms in both health and disease. As many biological species from evolution share protein orthologs, their mutual interactions can often be transferred, for example from an experiment in another organism to the corresponding pair of human proteins where the experiment has not yet been performed. Such correspondences can typically be revealed by text mining as researchers in one area often will not follow the literature in the other and *vice versa*.

We ran the text mining pipeline on the two full-text and two abstract corpora. In all cases we found that the AUC-value was far greater than 0.5, from which we conclude that the results were substantially better than random (Fig 2). The biggest gain in performance when using full-text was seen in finding associations between diseases and genes (Supplementary Table 2). Compared to MEDLINE, the traditional corpus used for biomedical text mining, there was an increase in the AUC from 0.85 to 0.91. The smallest gain was associations between proteins, which increased from 0.70 to 0.73. Likewise, the Core Full-texts always performed better than Core Abstracts, signifying that some associations are only reported in the main body of the text. Consequently, traditional text mining of abstracts will never be able to find this information.

**Fig 2:**
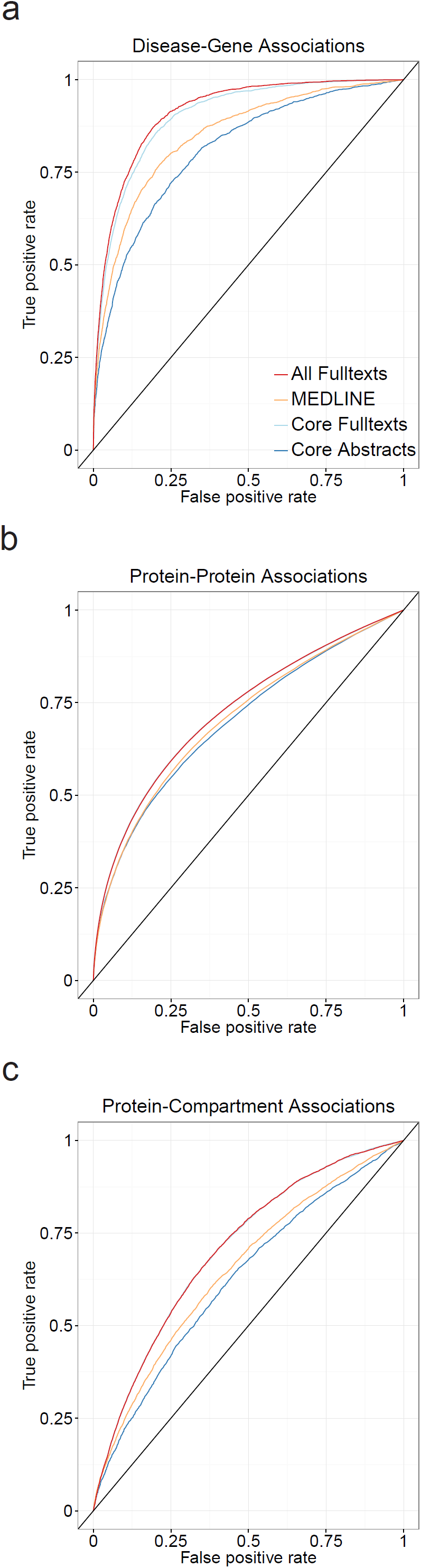
Benchmarking the four different corpora. In all cases the AUC is far greater than 0.5, indicating that the results obtained are better than random. The biggest gain in AUC is seen for disease-gene associations **(a)**, followed by protein-compartment associations **(c)** and protein-protein associations **(b)**.

It has previously been speculated if text mining of full-text articles may be more difficult and lead to an increased rate of false positives [3]. To investigate this we altered the weights of the scoring system. The scoring scheme used here has weights for within sentence, within paragraph and within document co-occurrences (see Methods). When setting the document weight to zero versus using the previously calibrated value we found that having a non-zero small value does indeed improve extraction of known facts in all cases (Supplementary Fig 4). Inspecting the gain in AUC we found that it is lower, compared to having a document weight (Supplementary Table 2). In one case, protein–protein associations, the MEDLINE abstract corpus outperforms the full-text articles. Abstracts are generally unaffected by the document weight, mainly because abstracts are almost always one paragraph. Overall, the difference in performance gain is largest for full-texts and lowest for abstracts and MEDLINE. Hence, all the full-text information is indeed valuable and necessary.

For practical applications, it is often necessary to have a low False Positive Rate (FPR). Accordingly, we evaluated the True Positive Rate (TPR) of the different corpora at the 10% FPR (TPR@10%FPR) (Fig 3). We found that full-texts have the highest TPR@10%FPR for disease-gene associations (Supplementary Table 3). When considering protein–protein associations and protein-compartment associations, full-texts perform equivalently to Core Abstracts and Core Full-texts. The result was similar to when we evaluated the AUC across the full range, removing the document weight has the biggest impact on the full-texts (Supplementary Fig 5), while abstracts remain unaffected.

**Fig 3:**
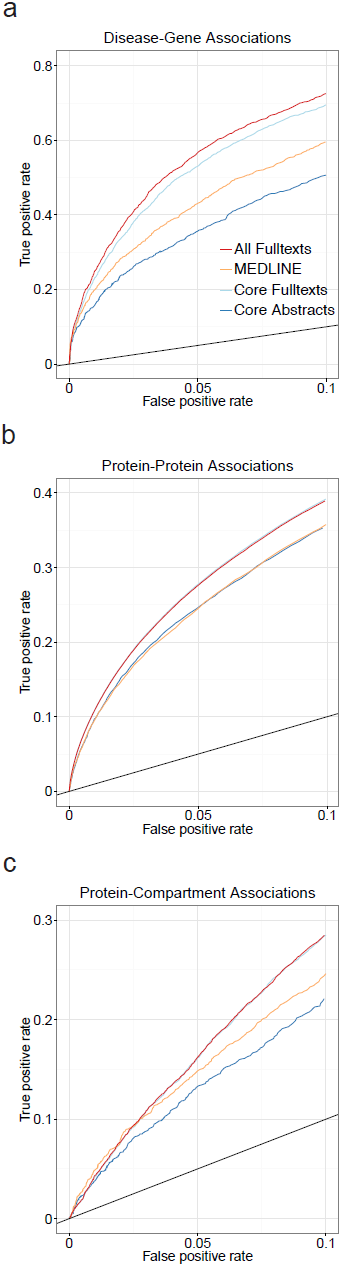
Benchmarking the four different corpora at low false positive rates. At a false positive rate of 10%, relevant to practical applications, the full-text corpus still outperforms the collection of MEDLINE abstracts for the extraction of disease-gene associations. Conversely, the performance is the same for protein-protein associations and protein-compartment associations.

## Discussion

We have investigated a unique corpus consisting of 15 million full-text articles and compared the results to the most commonly used corpus for biomedical text mining, MEDLINE. We found that the full-text corpus outperforms the MEDLINE abstracts in all benchmarked cases. To our knowledge, this is the largest comparative study to date of abstracts and full-text articles. We envision that the results presented here can be used in future applications for discovering novel associations from mining of full-text articles, and as a motivation to always include full-text articles when available and to improve the techniques used for this purpose.

The corpus consisted of 15,032,496 full-text documents, mainly in PDF format. 1,504,674 documents had to be discarded for technical reasons, primarily because they were not in English. Further, a large number of documents were also found to be duplicates or subsets of each other. On manual inspection we found that these were often conference proceedings, collections of articles etc., which were not easily separable without manual curation. We also managed to identify the list of references in the majority of the articles thereby reducing some repetition of knowledge that could otherwise lead to an increase in the false positive rate.

We have encountered and described a number of problems when working with full-text articles converted from PDF to TXT from a large corpus. However, the majority of the problems did not stem from the PDF to TXT conversion, which could potentially be solved using a layout aware conversion tool. Examples include LA-PDFText [27], SectLabel [26] of PDFX [25], of which the latter is not practical for very large corpora as it only exists as an online tool. Nonetheless, to make use of the large volume of existing articles it is necessary to solve these problems. Having all the articles in a structured XML format, such as the one provided by PubMed Central, would with no doubt produce a higher quality corpus. This may in turn further increase the benchmark results for full-text articles. Nevertheless, the reality is that many articles are not served that way. Consequently, the performance gain we report here should be viewed as a lower limit as we have sacrificed quality in favor of a larger volume of articles. The solutions we have outlined here will serve as a guideline and baseline for future studies.

The increasing article length may have different underlying causes, but one of the main contributors is most likely increased funding to science worldwide [28,29]. Experiments and protocols are consequently getting increasingly complex and interdisciplinary – aspects that also contribute to driving meaningful publication lengths upward. The increased complexity has also been found to affect the language of the articles, as it is becoming more specialized[30]. It was outside the scope of this paper to go further into socio-economic impact. We have limited this to presenting the trends from what could be computed from the meta-data.

Previous papers are – in terms of benchmarking – only making qualitative statements about the value of full-text articles as compared to text in abstracts. In one paper a single statement is made on the potential for extracting information, but no quantitative evidence is presented [31]. In a paper targeting pharmacogenomics it is similarly stated that that there are associations that only are found in the full-text, but no quantitative estimates are presented [20]. In a paper analyzing around 20,000 full-text papers a search for physical protein interactions was made, concluding that these contain considerable higher levels of interaction [19]. Again, no quantitative benchmarks were made comparing different sources. In this paper, we have made a detailed comparison of four different corpora that provides a strong basis for estimating the added value of using full-text articles in text mining workflows.

The results presented here are purely associational. Through rigorous benchmarking and comparison of a variety of biologically relevant associations, we have demonstrated that a substantial amount of relevant information is only found in the full body of text. Additionally, by modifying the document weight we found that it was important to take into account the whole document and not just individual paragraphs. Consequently, as text mining methods improve and become more sophisticated, the quantitative benchmarks will improve. Event-based text mining will be the next step for a deeper interpretation and extending the applicability of the results [5]. With more development it may also be possible to extract quantitative values, as has been demonstrated for pharmacokinetics [32]. However, this was outside the scope of this article.

The Named Entity Recognition (NER) system used depends heavily on the dictionaries and stop word lists. A NER system is also very sensitive to ambiguous words. To combat this we have used dictionaries from well-known and peer-reviewed databases, and we have included other dictionaries to avoid ambiguous terms. Other approaches to text mining have previously been extensively reviewed [10,14,32].

The full-text corpus presented here consists of articles from Springer, Elsevier and PubMed. However, we still believe that the results presented here are valid and can be generalized across publishers, to even bigger corpora. Preprocessing of corpora is an ongoing research project, and it can be difficult to weed out the rubbish when dealing with millions of documents. We have tried to use a process where we evaluate the quality of a subset of randomly selected articles repeatedly and manually, until it no longer improves.

## Methods

### MEDLINE Corpus

The MEDLINE corpus consists of 26,385,631 citations. We removed empty citations, corrections and duplicate PubMed IDs. For duplicate PubMed IDs we kept only the newest entry. This led to a total of 16,544,511 abstracts for text mining.

### PMC Corpus

The PubMed Central corpus comprises 1,488,927 freely available scientific articles (downloaded 27^th^ January 2017). Each article was retrieved in XML format. The XML file contains the article divided into paragraphs, article category and meta-information such as journal, year published, etc. Articles that had a category matching Addendum, Corrigendum, Erratum or Retraction were discarded. A total of 5,807 documents were discarded due to this, yielding a total of 1,483,120 articles for text mining. The article paragraphs were extracted for text mining. No further pre-processing of the text was done. The journals were categorized according to categories (described in the following section) by matching the ISSN number. The number of pages for each article was also extracted from the XML, if possible. Permission for use of the PMC corpus was obtained by the Technical Information Center of Denmark (DTU Library).

### TDM Corpus

The Technical Information Center of Denmark (DTU Library) TDM corpus is a collection of full-text articles from the publishers Springer and Elsevier, where the library has obtained permission for use in the context of text mining. The corpus covers the period from 1823 to 2016. The corpus comprises 3,335,400 and 11,697,096 full-text articles in PDF format, respectively. An XML file containing meta-data such as publication date, journal, etc. accompanies each full-text article. PDF to TXT conversion was done using pdftotext v0.47.0, part of the Poppler suite (poppler.freedesktop.org). 192 articles could not be converted to text due to errors in the PDF file. The article length, counted as the number of pages, was extracted from the XML file. If not recorded in the XML file we counted the number of pages in the PDF file using the Unix tool pdfinfo v0.26.5. Articles were grouped into four bins, determined from the 25%, 50%, and 75% quantiles, respectively. These were found to be 1-4 pages (0-25%), 5-7 pages (25-50%), 8-10 pages (50-75%) and 11+ pages (75%-100%). Each article was, based on the journal where it was published, assigned to one or more of the following seventeen categories: Health Sciences, Chemistry, Life Sciences, Engineering, Physics, Agriculture Sciences, Material Science and Metallurgy, Earth Sciences, Mathematical Sciences, Environmental Sciences, Information Technology, Social Sciences, Business and Economy and Management, Arts and Humanities, Law, Telecommunications Technology, Library and Information Sciences. Due to the large number of categories, we condensed anything not in the top-6 into the category “Other”. The top-six categories *health science, chemistry, life sciences, engineering, physics* and *agricultural sciences* make up 74.8% of the data (Supplementary Fig 2). The assignment of categories used in this study was taken from the existing index for the journal made by the librarians at the DTU Library. For the temporal statistics, the years 1823-1900 were condensed into one.

### Pre-processing of PDF-to-text converted documents

Following the PDF-to-text conversion of the Springer and Elsevier articles we ran a language detection algorithm implemented in the python package langdetect v1.0.7 (https://pypi.python.org/pypi/langdetect). We discarded 902,415 articles that were not identified as English. We pre-processed the remaining raw text from the articles as follows:

1. Non-printable characters were removed using the POSIX filter [[:^print:]].
2. A line of text was removed if digits make up more than 10% of the text, or symbols make up more than 10% of the text, or lowercase text was less than 50%. Symbols are anything not matching [0-9A-Za-z].
3. Removal of acknowledgements and reference-or bibliography-lists using a rule-based system explained below.
4. Text was split into sentences and paragraphs using a rule-based system described below.

We assumed that acknowledgements and reference lists are always at the end of the article. Upon encountering either of the terms: “acknowledgement”, “bibliography”, “literature cited”, “literature”, “references”, and the following misspellings thereof: “refirences”, “literatur”, “références”, “referesces”. In some cases the articles had no heading indicating the start of a bibliography. We tried to take these cases into account by constructing a RegEx that matches the typical way of listing references (e.g. [1] Westergaard, …). Such a pattern can be matched by the RegEx “^\[\d+\]\s[A-Za-z]”. The other commonly used pattern, “1. Westergaard, …”, was avoided since it may also indicate a new heading. Keywords were identified based on several rounds of manual inspection. In each round, 100 articles in which the reference list had not been found was randomly selected and inspected. We were unable to find references in 286,287 and 2,896,144 Springer and Elsevier articles, respectively. Manual inspection of 100 randomly selected articles revealed that these articles indeed did not have a reference list or that the pattern was not easily describable with simple metrics, such as keywords and RegEx. Articles without references were not discarded.

The PDF to text conversion often breaks up paragraphs and sentences, due to new page, new column, etc. Paragraph and sentence splitting was performed using a ruled-based system. If the previous line of text does not end with a “.!?”, and the current line does not start with a lower-case letter, it is assumed that the line is part of the previous sentence. Otherwise, the line of text is assumed to be a new paragraph.

### Text article filtering

A number of Springer and Elsevier documents were removed due to technical issues post pre-processing. An article was removed if:

1. Article contained no text post-preprocessing (51,399 documents).
2. Average word length was below the 2% quantile (263,902 documents).
3. Article contained specific keywords, described below (286,958 documents).

Some PDF files without texts are scans of the original article (point 1). We did not attempt to make an optical character recognition conversion (OCR) as the old typesetting fonts often are less compatible with present day OCR programs, and this can lead to text recognition errors [33,34]. For any discarded document, we still used the meta-data to calculate summary statistics. In some cases the PDF to text conversion failed, and produced non-sense data with a white space between the characters of a majority of the words (point 2). To empirically determine a cutoff we gradually increased the cutoff and repeatedly inspected 100 randomly selected articles. At the 2% quantile we saw no evidence of broken text.

Articles with the following keywords in the article were discarded: Author Index, Key Word Index, Erratum, Editorial Board, Corrigendum, Announcement, Books received, Product news, and Business news (point 3). These keywords were found as part of the process of identifying acknowledgements and reference lists. Further, any article that was available through PubMed Central was preferentially selected by matching doi identifiers. This left a total of 14,549,483 full-text articles for further analysis.

Some articles were not separable, or were subsets of others. For instance, conference proceedings may contain many individual articles in the same PDF. We found 1,911,365 articles in which this was the case. In these cases we removed the duplicates, or the shorter texts, but kept one copy for text mining. In total, we removed 898,048 duplicate text files.

The majority of articles had a separate abstract. We matched articles from PubMed Central to their respective MEDLINE abstract using the PMCID to PubMed ID conversion file available from PMC. Articles from Springer and Elsevier typically had a separate abstract in the meta-data. Any abstract from an article that was part of the 1,911,365 articles that could not be separated was removed. This led to a total of 10,376,626 abstracts for which the corresponding full-text was also included downstream, facilitating a comparative analysis.

### Text mining of articles

We performed text mining of the articles using a Named Entity Recognition (NER) system, described earlier[35–38]. The software is open source and can be downloaded from https://bitbucket.org/larsjuhljensen/tagger. The NER approach is dictionary based, and thus depends on well-constructed dictionaries and stop word lists. We used the gene names from the STRING dictionary v10.0 [35], disease names from the Disease Ontology (DO) [39] and compartment names from the Gene Ontology branch cellular component [40]. Stop word lists were all created and maintained in-house. Pure NER based approaches often struggles with ambiguity of words. Therefore, we included additional dictionaries that we do not report the results from. If any identified term was found in multiple dictionaries, it was discarded due to ambiguity. The additional dictionaries include small molecule names from STITCH [41], tissue names from the Brenda Tissue Ontology [42], Gene Ontology biological process and molecular function [40], and the mammalian phenotype ontology [43]. The latter is a modified version made to avoid clashes with the disease ontology. The dictionaries can be downloaded from http://download.jensenlab.org/.

In the cases where the dictionary was constructed from an ontology co-occurrences were backtracked through all parents. E.g. the term type 1 diabetes mellitus from the Disease Ontology is backtracked to its parent, diabetes mellitus, then to glucose metabolism disease, etc.

Co-occurrences were scored using the scoring system described in [44]. In short, a weighted count for each pair of entities (e.g. disease-gene) was calculated using the formula,

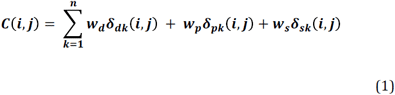

where *δ* is an indicator function taking into account whether the terms *i,j* co-occur within the same document (d), paragraph (p), or sentence (s). *w* is the co-occurrence weight here set to 1.0, 2.0, and 0.2, respectively. Based on the weighted count, the score *S(i,j)* was calculated as,

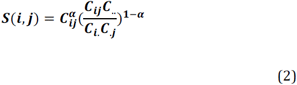

where *α* is set to 0.6. All weights were optimized using the KEGG pathway maps as benchmark (described further below). The *S* scores were converted to *Z* scores, as described earlier [45].

### Benchmarking of associations

PPIs were benchmarked using pathway maps from the KEGG database [46]. Any two proteins in the same pathway were set to be a positive example, and any two proteins present in at least one pathway, but not the same, were set as a negative example. This approach assumes that the pathways are near complete and includes all relevant proteins. The same approach has been used for the STRING database [44]. The disease–gene benchmarking set was created by setting the disease-gene associations from UniProt [47] and Genetics Home Reference (https://ghr.nlm.nih.gov/, accessed 23^th^ March 2017) as positive examples. The positive examples were then shuffled, and the shuffled examples were set as negative examples. Shuffled examples that ended up overlapping with the positive examples were discarded. This approach has previously been described [36]. The protein–compartment benchmark set was created by extracting the compartment information for each protein from UniProt and counting these as positive examples. For every protein found in at least one compartment, all compartments where it was not found were set as negative examples. The same approach has been used previously [38].

Receiver Operating Characteristic (ROC) curves were created by gradually increasing the *Z*-score and calculating the True Positive Rate (TPR) and False Positive Rate (FPR), as described in eqs. (3) and (4).

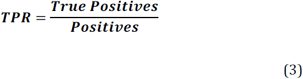

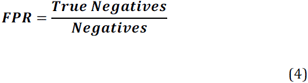

We compare the ROC curves by the Area Under the Curve (AUC), a metric ranging from 0 to 1.

## Acknowlegements

We would like to acknowledge funding from ActionableBiomarkersDK, a grant from DeIC, the Danish e-Infrastructure Collaboration, as well as the Novo Nordisk Foundation (grant agreement NNF14CC0001).

## Supporting Information Captions

**S1 Fig:**
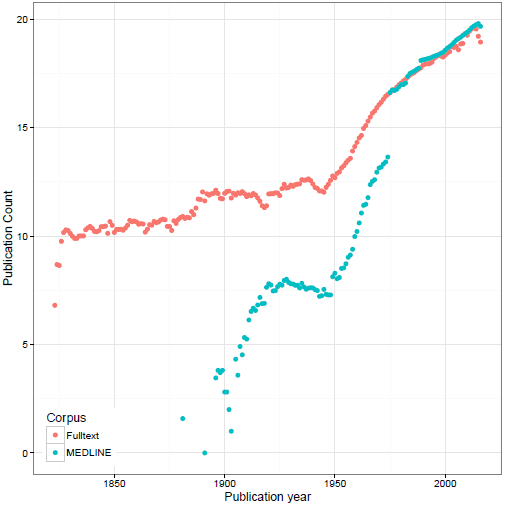
Number of publications per year on the log scale.

**S2 Fig:**
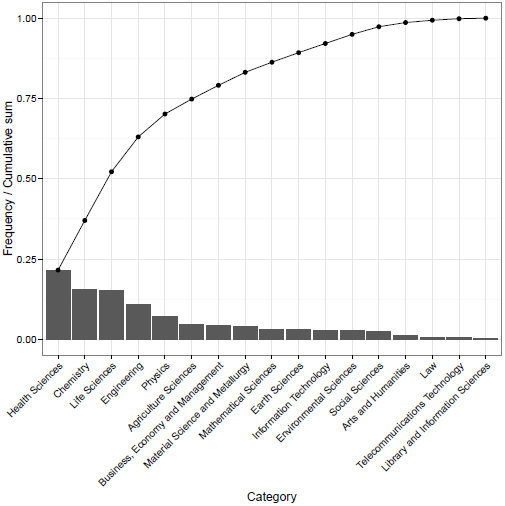
Category overview across all journals and years. The bar chart indicates the frequency, whilst the line is the cumulative sum. The first six categories contribute 74,8%. Due to the large number of categories, anything outside the top-6 was condensed into the joint category “Other”.

**S3 Fig:**
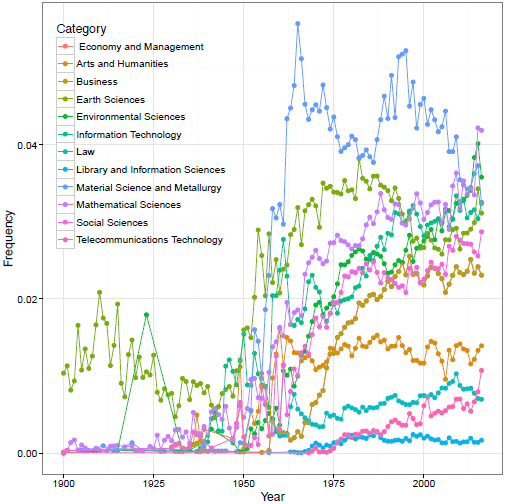
Temporal trend for the categories embedded in the “Other” category. We note that the category has grown as a whole, but that the growth is not tied to one category.

**S4 Fig:**
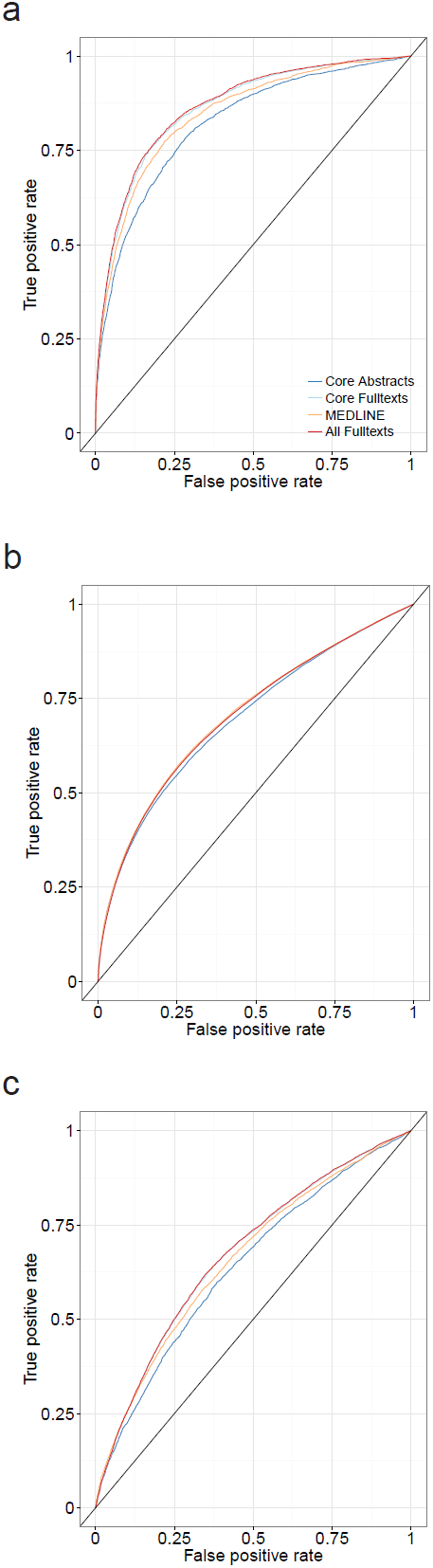
Benchmarking the four different corpora not using a document weight. (a-c) The increase in performance has fallen, compared to including a document weight. In one case, protein-protein associations, the MEDLINE corpus outperforms the full text articles.

**S5 Fig:**
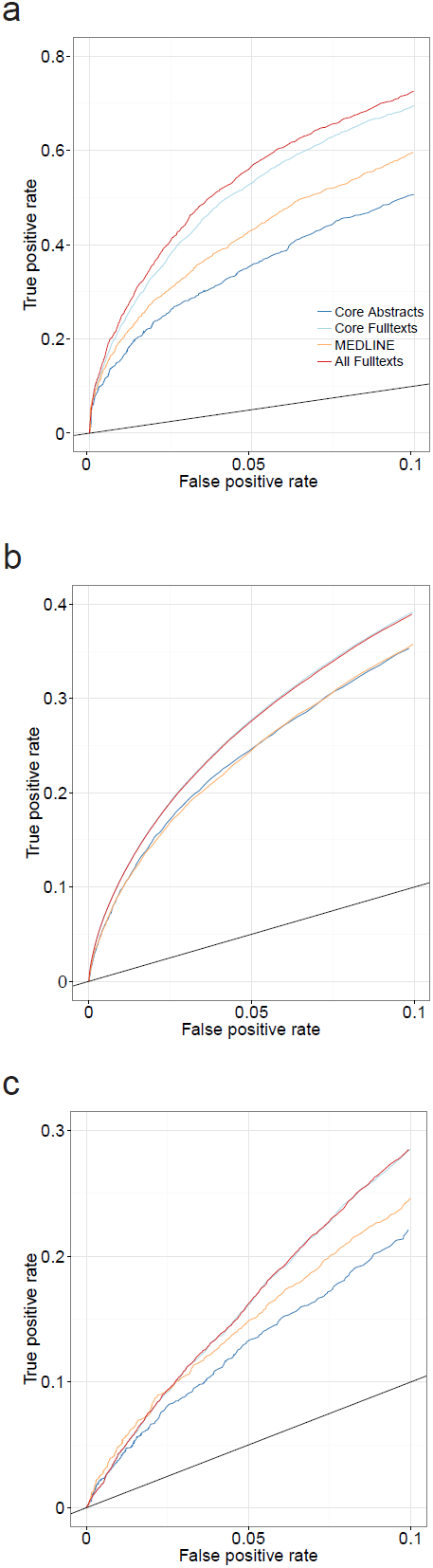
Benchmarking the four different corpora at a low false positive rate not using a document weight. The increase in performance has fallen. In one case, for protein-protein associations, the MEDLINE corpus outperforms the full text articles.

**S1 Table:** The top 15 journals in the corpora.

**S2 Table:** Area under the curve (AUC) for the four different corpora, with and without document weight for scoring co-occurrences.

**S3 Table:** True Positive Rate at 10% False Positive Rate (TPR@10%FPR) for the four different corpora, with and without document weight for scoring co-occurrences.

